# Rats exhibit similar biases in foraging and intertemporal choice tasks

**DOI:** 10.1101/497321

**Authors:** Gary A. Kane, Aaron M. Bornstein, Amitai Shenhav, Robert C. Wilson, Nathaniel D. Daw, Jonathan D. Cohen

## Abstract

Animals, including humans, consistently exhibit myopia in two different contexts: foraging, in which they harvest locally beyond what is predicted by optimal foraging theory, and intertemporal choice, in which they exhibit a preference for immediate vs. delayed rewards beyond what is predicted by rational (exponential) discounting. Despite the similarity in behavior between these two contexts, previous efforts to reconcile these observations in terms of a consistent pattern of time preferences have failed. Here, via extensive behavioral testing and quantitative modeling, we show that rats exhibit similar time preferences in both contexts: they prefer immediate vs. delayed rewards and they are sensitive to opportunity costs — delays to future decisions. Further, a quasi-hyperbolic discounting model, a form of hyperbolic discounting with separate components for short-and long-term rewards, explains individual rats’ time preferences across both contexts, providing evidence for a common mechanism for myopic behavior in foraging and intertemporal choice.

## Introduction

Serial stay-or-search problems are ubiquitous across many domains, including employment, internet search, mate search, and animal foraging. For instance, in patch foraging problems, animals must choose between an immediately available opportunity for reward or the pursuits of potentially better but more distal opportunities. The optimal behavior in foraging tasks, described by the Marginal Value Theorem (MVT; Charnov 1976), is to maximize long-term average reward rate by choosing the immediately available opportunity if it provides a reward rate greater than the average reward rate across all alternative options, which includes the costs of accessing those options. Animals tend to follow the basic predictions of long-term reward maximization: they are generally more likely to pursue opportunities for larger vs. smaller rewards and, if the costs of searching for alternatives is greater, they are more likely to pursue opportunities for smaller rewards (Constantino and Daw, 2015; Hayden et al., 2011; Kane et al., 2017; Stephens and Krebs, 1986).

Although animal behavior follows the basic predictions of optimal foraging behavior described by MVT, in the majority of studies across a variety of species, including humans, non-human primates, and rodents, animals exhibit a consistent bias towards pursuing immediately available rewards relative to predictions of MVT, often referred to as “overharvesting” (Carter and Redish, 2016; Constantino and Daw, 2015; Hayden et al., 2011; Kane et al., 2017; Kolling et al., 2012; Nonacs, 2001; Shenhav et al., 2014; Wikenheiser et al., 2013). Prior studies have proposed two explanations for overharvesting: subjective costs, such as an aversion to rejecting an immediately available reward (Carter and Redish, 2016; Wikenheiser et al., 2013); and diminishing marginal returns, by which larger rewards are not perceived as proportionally larger than smaller rewards (Constantino and Daw, 2015). But these hypotheses have never been systematically compared in a set of experiments designed to directly test their predictions. Furthermore, according to these hypotheses, the perceived value of rewards does not change if delays occur before or after reward is received, so long as they occur before future decisions (e.g. an inter-trial interval). In this respect, the predictions made by these hypotheses are not compatible with an otherwise seemingly similar bias that is widely observed in standard intertemporal choice tasks (also referred to as delay discounting or self-control tasks): a preference for smaller, more immediate rewards over larger, delayed rewards (Ainslie, 1992; Kirby, 1997).

The preference for more immediate rewards in intertemporal choice tasks is commonly explained in one of two ways: temporal discounting or short-term rate maximization. According to temporal discounting, the perceived value of a future reward is discounted by the time until its receipt. Discounting can be adaptive in unstable environments if the environment is likely to change before future rewards can be acquired; in such cases, it is appropriate to place greater value on more predictable rewards available in the near future. Under this hypothesis, the normative discount function is exponential, since the rate of discounting remains constant over time (Gallistel and Gibbon, 2000; Kacelnik and Todd, 1992). However, animal preferences typically follow a hyperbolic-like form: the rate of discounting is steeper initially and decreases over time (Gallistel and Gibbon, 2000; Kacelnik and Todd, 1992; Thaler, 1981). This yields inconsistent time preferences or preference reversals: an animal may prefer to wait longer for a larger reward if both options are distant, but will change their mind and prefer the smaller reward as the time to both options draws near (Ainslie, 1992; Gallistel and Gibbon, 2000; Kacelnik and Todd, 1992; Kirby, 1997). Short-term rate maximization rules predict that animals attempt to maximize reward on a shorter timescale (i.e. local reward rate), rather than maximizing all future rewards (i.e. global reward rate). One advantage to maximizing local reward rates is that animals may be able to better estimate the value of rewards in the short-term, thus making better decisions over this time than if they considered all future reward (Gabaix and Laibson, 2017; Stephens, 2002; Stephens et al., 2004). In support of this hypothesis, in intertemporal choice tasks, animals typically attend only to delays between decisions and receiving rewards and they are insensitive to post-reward delays (i.e., delays between receiving reward and making the next decision; Bateson and Kacelnik 1996; Blanchard et al. 2013; Pearson et al. 2010; Stephens and Anderson 2001).

Despite the similarities between overharvesting and the preference for more immediate rewards in intertemporal choice tasks, prior attempts to use temporal discounting and/or short-term rate maximization functions fit to intertemporal choice data to predict foraging behavior have failed (Blanchard and Hayden, 2015; Carter et al., 2015; Carter and Redish, 2016). There are two possible explanations for why intertemporal choice models have failed to predict foraging behavior: i) differences in the decision horizon of intertemporal choice and foraging tasks, and ii) the choice of discounting function. i) Models of intertemporal choice tasks usually consider rewards for the current trial and not rewards on future trials since, in these tasks, reward opportunities on future trials are often independent of the current decision. This is not true of foraging tasks, in which future opportunities for rewards depend on the current decision. Thus, this difference in decision horizon may make it difficult to explain foraging data using discounting models fit to intertemporal choice data. ii) Additionally, these studies have only examined standard, single-parameter exponential and hyperbolic discounting functions. More flexible forms of temporal discounting, such as quasi-hyperbolic discounting, which has separate terms for short-and long-term rewards that correlate with activity in limbic and fronto-parietal networks respectively (Laibson, 1997; McClure et al., 2007, 2004), have never been tested in these contexts.

In the present study, we found that rats exhibit similar time preferences in foraging and intertemporal choice tasks and that time preferences in both tasks can be explained by a quasi-hyperbolic discounting model that, in both contexts, considers future rewards. Rats were tested in a series of patch foraging tasks and an intertemporal choice task. In foraging tasks, they followed the basic predictions of long-term rate maximization: they stayed longer in patches that yielded greater rewards and when the cost of searching was greater. But under certain conditions, they violated these predictions in a manner consistent with time preferences: they stayed longer in patches when given larger rewards with proportionately longer delays, and they exhibited greater sensitivity to pre- vs. post-reward delays. Similarly, in an intertemporal choice task, rats exhibited greater sensitivity to pre- vs. post-reward delays. Using these data, we tested several models to determine if temporal discounting or biases in time perception based on short-term rate maximization, such as ignoring post-reward delays, could explain rats’ behavior across tasks. One model, a quasi-hyperbolic discounting model (Laibson, 1997; McClure et al., 2007), provided the best fit to rat behavior across all experiments. Furthermore, the quasi-hyperbolic discounting model proved to be externally valid: discounting functions fit to foraging data provided as good a fit to intertemporal choice data as discounting functions fit directly to intertemporal choice data for the majority of rats. These findings suggest that rats exhibit similar biases in the two tasks, and quasi-hyperbolic discounting may be a common mechanism for suboptimal decision-making across tasks.

## Results

### Rats consider long-term rewards, but exhibit a bias in processing pre- vs. post-reward delays

Long Evans rats (n = 8) were tested in a series of patch foraging tasks in operant conditioning chambers (Kane et al., 2017). To harvest reward (10% sucrose water) from a patch, rats pressed a lever on one side of the front of the chamber (left or right) and reward was delivered in an adjacent port. After a post-reward delay (inter-trial interval or ITI), rats again chose to harvest a smaller reward or to leave the patch by nose poking in the back of the chamber. A nose poke to leave the patch caused the harvest lever to retract and initiated a delay to control the time to travel to the next patch. After the delay, the opposite lever extended (e.g. if the left lever was extended previously, the right lever would be extended now), and rats could then harvest from (or leave) this replenished patch (Fig. S1).

In four separate experiments, we manipulated different variables of the foraging environment: i) in the “Travel Time Experiment,” a 10 s vs. 30 s delay was imposed between patches, ii) in the “Depletion Rate Experiment,” reward depleted at a rate of 8 vs. 16 µL per harvest, iii) in the “Scale Experiment,” the overall magnitude of rewards and delays was varied, such that in one condition, the size of rewards and length of delays was twice that of the other. iv) Finally, in the “Pre-vs-Post Experiment,” the placement of delays was varied, such that the total time to harvest reward remained constant, but in one condition there was no pre-reward delay and ∼13 s post reward delay, and in the other there was a 3 s pre-reward delay and ∼10 s post-reward delay. Parameters for each experiment are shown in Table 1. For each condition within each experiment, rats were trained for 5 days and tested for an additional 5 days; all behavioral data presented is from the 5 test days. The order of conditions within each experiment was counterbalanced across rats. Every patch visit was included for analysis; mixed effects models were used to examine the effect of task condition on the number of trials spent in each patch, with random intercepts and random slopes for the effect of task condition across rats. To compare rat behavior to the optimal behavior in each condition, a mixed effects model was used to test the effect of task condition on the difference between the number of trials spent in each patch and the optimal number of trials for that patch, with random intercepts and slopes for each rat. For this mixed effects model, an intercept of zero indicates optimal performance, and the slope indicates the change in behavior relative to the optimal behavior between conditions.

**Table 1.** Parameters for each of the first 4 foraging experiments. Harvest time = time to make a decision to harvest + pre-reward delay + inter-trial interval. To control reward rate in the patch, the inter-trial interval was adjusted relative to the decision time to hold the harvest time constant.

| Experiment | Condition | Start Reward | Depletion Rate | Pre-Reward Delay | Harvest Time | Travel Time |
| --- | --- | --- | --- | --- | --- | --- |
| travel time | 10s<br>30 s | 60, 90, or 120 μL | -8 μL | 0 s | 10 s | 10 s<br>30 s |
| depletion rate | -8 μL<br>-16 μL | 90 μL | -8 μL<br>-16 μL | 0 s | 12 s | 12 s |
| scale | 90 μL/10 s<br>180 μL/20 s | 90 μL<br>180 μL | -8 μL<br>-16 μL | 0 s | 10 s<br>20 s | 10 s<br>20 s |
| pre-vs-post | 0 s<br>3 s | 90 μL | -8 μL | 0 s<br>3 s | 15 s | 15 s |

The Travel Time Experiment was designed to test the two main predictions of MVT: i) that animals should stay longer in patches that yield greater rewards and ii) animals should stay longer in all patches when the cost of traveling to a new patch is greater. In this experiment, rats encountered three different patch types within sessions, which started with varying amount of reward (60, 90, or 120 µL) and depleted at the same rate (8 µL/harvest). The delay between patches was either 10 s or 30 s; each travel time delay was tested in its own block of sessions and the order was counterbalanced across rats, with a range of 87-236 patches visited per condition per rat. As predicted by MVT, rats stayed for more trials in patch types that started with larger reward volume (β = 118.091, SE = 1.862, t(2490.265) = 63.423, p <.001), indicating that rats considered reward across future patches. Rats also stayed longer in all patch types when time between patches was longer (β = 1.893, SE =.313, t(118.839) = 6.040, p <.001; Fig. 1A), indicating sensitivity to opportunity costs. However, rats uniformly overharvested relative to predictions of MVT (β_rat-MVT_ = 3.396, SE =.176, t(6.960) = 19.269, p <.001). The degree to which rats overharvested was not significantly different between the 10 s and 30 s travel conditions (β_10 s-30 s_ =.304, SE =.155, t(7.3857) = 1.964, p =.088).

**Figure 1.**
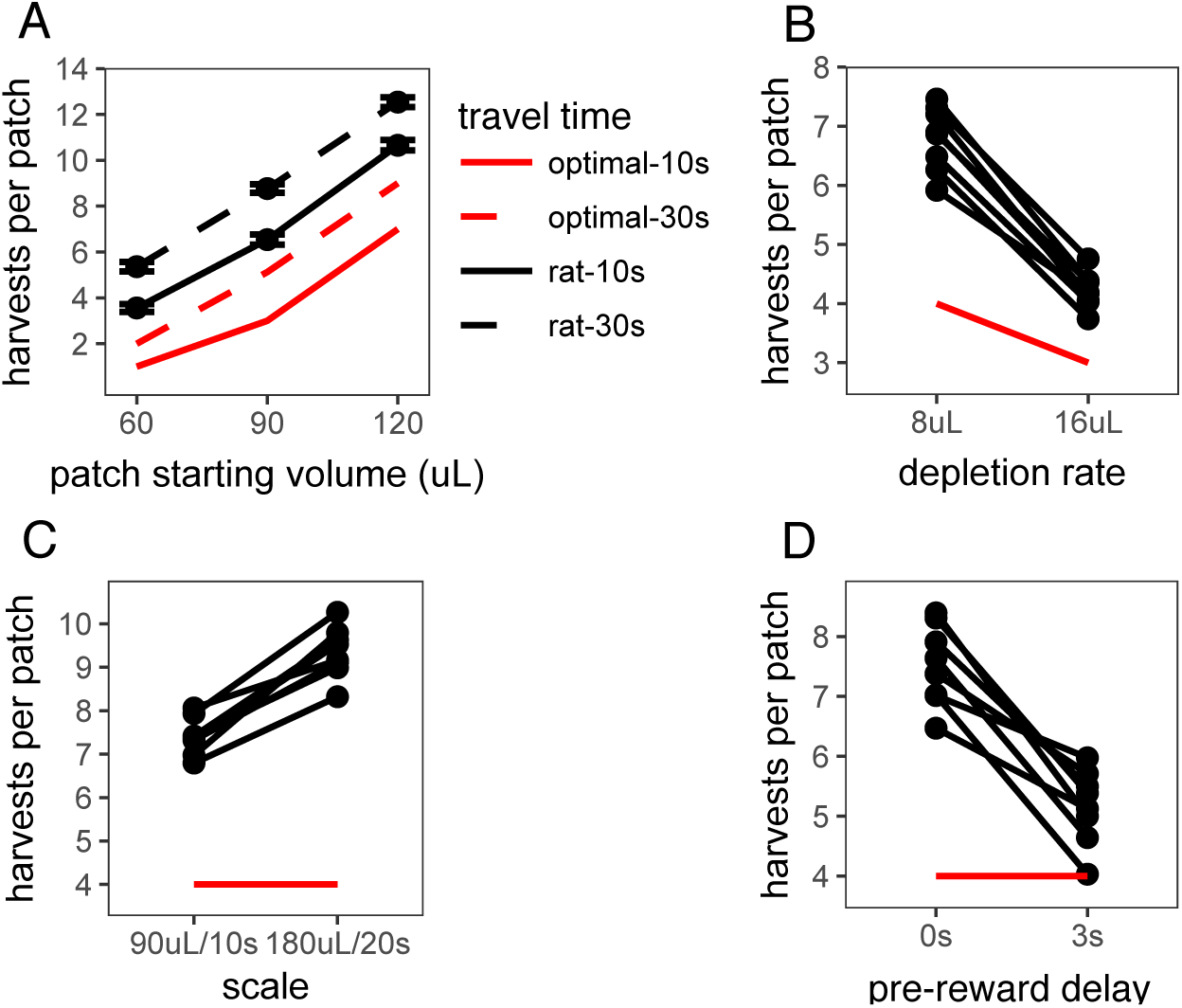
Rat foraging behavior in the A) Travel Time, B) Depletion Rate, C) Scale, and D) Pre-vs-Post Experiments. In A, points and error bars represent mean ± standard error. In B-D, points and connecting lines represent behavior of each individual rat. Red lines indicate optimal behavior (per MVT).

The Depletion Rate Experiment tested another critical variable in foraging environments: the rate of reward depletion within a patch. Quicker reward depletion causes the local reward rate to deplete to the long-run average reward rate quicker, thus MVT predicts earlier patch leaving. Within sessions, rats encountered a single patch type (starting volume of 90 µL) that depleted at a rate of either 8 or 16 µL/trial, tested in separate sessions and counterbalanced, with a range of 152-283 patches visited per condition per rat. As predicted by MVT, rats left patches earlier when patches depleted more quickly (β = 2.589 harvests, SE =.155, t(7.000) = 16.75, p <.001; Fig. 1B). But, again, rats stayed in patches longer than is predicted by MVT (β_rat-MVT_ = 2.005, SE =.134, t(7.004) = 14.97, p <.001). Rats overharvested to a greater degree in the 8 µL depletion condition than the 16 µL depletion condition (β_8 µL-16 µL_ = 1.589, SE =.155, t(7.000) = 10.28, p < .001).

These first two experiments confirm that rats qualitatively follow the predictions of MVT, but consistently overharvest. There are many possible explanations for this pattern of overharvesting, including an aversion to leaving the offer of reward within a patch and diminishing marginal returns (Carter and Redish, 2016; Constantino and Daw, 2015; Wikenheiser et al., 2013). The Scale Experiment was conducted in an effort to distinguish between these hypotheses by manipulating the scale of time delays and rewards. Long-term rate maximization predicts that an increase in reward size in proportion to reward delay should have no effect on the number of harvests per patch, as the reward rate across trials would be equal. But if animals’ perception of reward or time is nonlinear, a manipulation of scale will affect their subjective point of equality and predict a change in behavior across the two environments. The scale of rewards and delays was manipulated in the following manner: patches started with (A) 90 or (B) 180 µL of reward, depleted at a rate of (A) 8 or (B) 16 µL/trial, and the duration of harvest trials and travel time between patches was (A) 10 or (B) 20 s. Rats visited a range of 60-212 patches per condition. They overharvested in both A and B conditions (β_rat-MVT_ = 4.374, SE =.153, t(6.900) = 28.597, p <.001) and, contrary to predictions of MVT, they stayed in patches significantly longer and overharvested to a greater degree in the B condition that provided larger rewards but at proportionately longer delays (β = 1.937, SE =.193, t(6.972) = 9.996, p <.001; Fig. 1C). This finding suggests that a nonlinearity in the perception of reward value and/or time contributes to overharvesting. However, there are still other possible explanations for this behavior, including diminishing marginal returns for larger rewards, biases in time perception such as insensitivity to post-reward delays, and/or temporal discounting.

To distinguish between biases in perception of reward, such as diminishing marginal returns, and time, such as temporal discounting or insensitivity to post-reward delays, the Pre-vs-Post Experiment directly tested rats sensitivity to time delays before vs. after reward. In this experiment, in one condition, rats received reward immediately after lever press followed by a post-reward delay of ∼13 s before the start of the next trial. In the other condition, there was a 3 s pre-reward delay between lever press and receiving reward followed by a shorter post-reward delay of ∼10 s. The total time of each trial was held constant between conditions (15 s total), so there was no difference in reward rates. Both MVT and diminishing marginal returns predict that the placement of delays is inconsequential and that rats will behave similarly in both conditions. Both temporal discounting and insensitivity to post-reward delays predict that rats will value the immediate reward more than the delayed reward and thus, would leave patches earlier in the condition with the pre-reward delay. Consistent with predictions of temporal discounting and insensitivity to post-reward delays, and contrary to predictions of MVT and diminishing marginal returns, rats left patches earlier in the environment with the pre-reward delay (β = 2.345, SE =.313, t(7.017) = 7.503, p <.001; Fig. 1D). This result suggests that a bias in rats’ perception of time or the way in which they perceive reward values over time contributes to overharvesting.

To determine whether the preference for immediate rewards can be explained by insensitivity to post-reward delays, a fifth foraging experiment, the “Post-Reward Delay Experiment,” directly tested rats’ sensitivity to post-reward delays. A separate cohort of rats (n = 8) was used for this experiment. Rats were tested in two conditions in this experiment: a short (3 s) or long (12 s) post-reward delay. The total time of harvest trials was not held constant; the longer post-reward delay increased the time to harvest from the patch. Since the longer post-reward delay increases the cost of harvesting from the patch relative to the cost of travelling to a new patch, MVT predicts that rats should leave patches earlier. Prior studies of intertemporal choice behavior suggest that animals are insensitive to post-reward delays, and that they are only concerned with maximizing short-term reward rate (Bateson and Kacelnik, 1996; Blanchard et al., 2013; Stephens and Anderson, 2001). A common formalization of this hypothesis assumes that reward rate is maximized only over the time to receive the next reward (e.g. short-term rate = next reward / delay to next reward; Bateson and Kacelnik 1996; Carter and Redish 2016; Stephens and Anderson 2001). This form of short-term rate maximization would predict that the increase in post-reward delay should have no effect on rat behavior. As predicted by MVT, but not this form of short-term rate maximization, rats were sensitive to the post-reward delay, leaving patches earlier in the 12 s delay condition (β = 1.411 trials, SE =.254, t(6.966) = 5.546, p <.001; Fig. 2A). As in other experiments, rats overharvested (β_rat-MVT_ = 3.332 trials, SE =.285, t(7.041) = 11.704, p <.001), and there was no difference in the degree to which rats overharvested between the 3 s and 12 s delay conditions (β_3 s-12 s_ =.340 trials, SE =.286, t(6.963) = 1.188, p =.274). These data suggest that rats consider the timing and magnitude of rewards beyond the next expected reward when making decisions, and that earlier patch leaving in the Pre-vs-Post Experiment is likely due to hypersensitivity to pre-reward delays and not insensitivity to post-reward delays.

**Figure 2.**
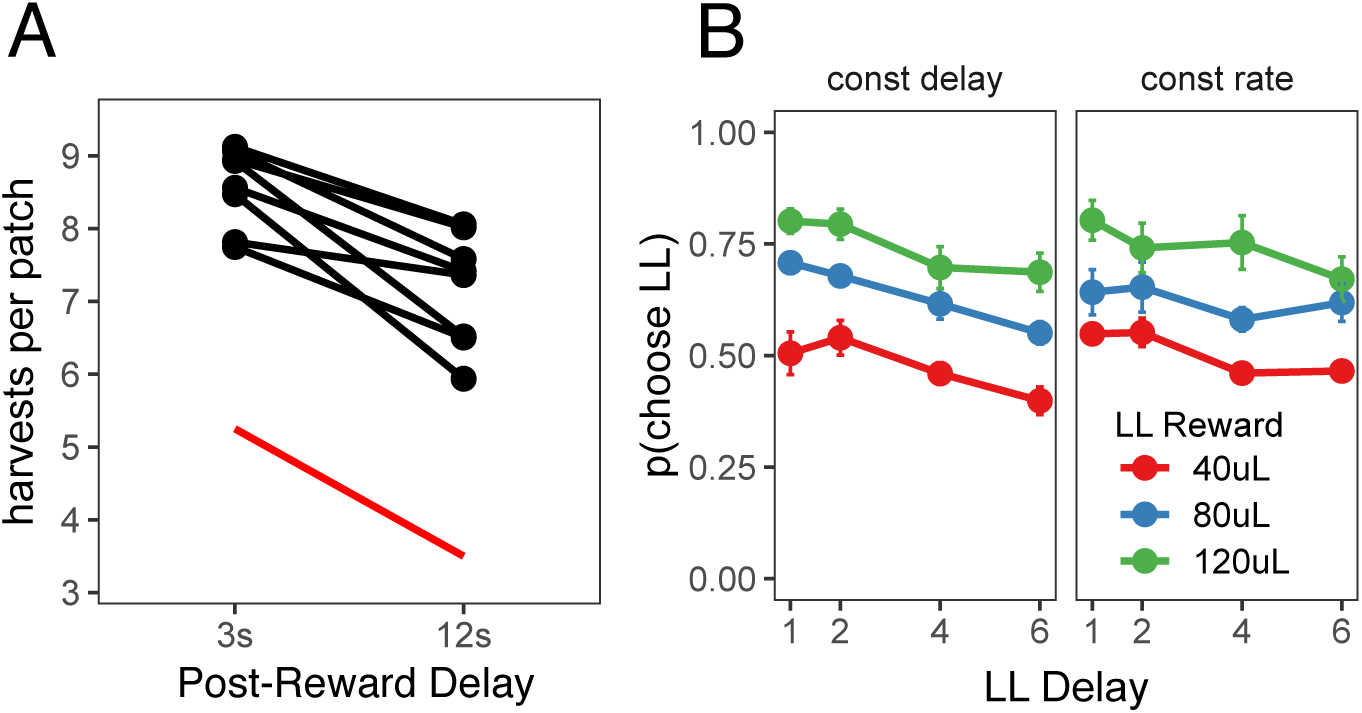
A) Rat behavior in the Post-Reward Delay Experiment. Points and lines represent behavior of individual rats. Red line indicates optimal behavior (per MVT). B) Rat behavior in the two-alternative intertemporal choice task. Points and error bars represent mean ± standard error for each condition.

The data from the foraging experiments described above suggest that rats exhibit time preferences in the foraging task. In a final “Intertemporal Choice Experiment,” we tested whether the same rats that participated in the Post-Reward Delay Experiment would exhibit similar time preferences in a standard intertemporal choice (i.e. a delay-discounting) task. This task consisted of a series of 20-trial episodes. On each trial, rats pressed either the left or right lever to receive a smaller-sooner (SS) reward of 40 µL after a 1 s delay or a larger-later (LL) reward of 40, 80, or 120 µL after a 1, 2, 4, or 6 s delay. For the first 10 trials of each episode, rats were forced to press either the left or right lever to learn the value and delay associated with that lever (only one lever extended on each of these trials). For the last 10 trials of an episode, both levers extended and rats were free to choose. The LL reward value and delay, and the LL lever (left or right) were randomly selected at the start of each episode. Rats were tested in two different versions of this task: one in which the post-reward delay was held constant, such that the longer pre-reward delays reduced reward rate (constant delay); and another in which the time of the trial was held constant, such that longer pre-reward delays resulted in shorter post-reward delays to keep reward rate constant (constant rate). MVT, which maximizes long-term reward rate, predicts that rats would be sensitive to the pre-reward delay in the constant delay condition but not the constant trial condition (in which the pre-reward delay does not affect reward rate).

Rats were given 3 training sessions to learn the structure of the intertemporal choice task after previously being tested in the foraging task, then they were tested for an additional 13 sessions in each condition, participating in a range of 590-2810 free choice trials per condition (constant delay vs. constant rate). Each free choice trial within each episode was counted as a separate observation. Choice data were analyzed using a generalized linear mixed-effects model (i.e. a mixed-effects logistic regression) to examine the effect of the size of the LL reward, the length of the LL delay, task condition (constant delay vs. constant rate), and their interactions on decisions to choose the LL vs. SS option, with random intercepts and random slopes for the effects of LL reward, LL delay, and task condition for each rat. Three post-hoc comparisons were used to test the effects of i) LL reward and ii) LL delay within each condition, and iii) LL delay between the constant delay and constant rate conditions (Fig. 2B). i) In both conditions, rats were more likely to choose larger LL rewards (constant delay: β =.477, SE =.090, χ^2^(1) = 28.320, p <.001; constant rate: β =.450, SE =.089, χ^2^(1) = 25.378, p <.001), showing that they were sensitive to reward magnitude. ii) They were also sensitive to the pre-reward delay in both conditions (constant delay: β = −.240, SE =.023, χ^2^(1) = 104.882, p <.001; constant rate: β = −.152, SE =.022, χ^2^(1) = 46.919, p <.001). On average, rats were equally likely to select the LL option across conditions — the main effect of task condition was not significant (β =.010, SE =.105, z =.092, p =.927). iii) However, rats were less sensitive to increasing pre-reward delays when pre-reward delays did not affect reward rate (the constant rate condition), indicated by a change in LL delay slope between conditions (β =.088, SE =.026, χ^2^(1) = 11.376, p <.001). Overall, rats exhibited similar time preferences in the foraging and intertemporal choice tasks: they valued rewards less with longer delays until receipt but they were sensitive to opportunity costs (e.g. time delays between receiving reward and future decisions).

### Quasi-hyperbolic discounting best explains behavior across all tasks

To test whether a common set of cognitive biases could explain time preferences in both the foraging and intertemporal choice tasks, both tasks were modeled as continuous time semi-markov processes. States were represented as the time between each event in each of the tasks (e.g. cues turning on/off, lever press, reward delivery; for state space diagrams of both tasks, see Fig. S2). The value of a given state was the discounted value of all future rewards available from that state, and the agent chose the option that yielded the greatest discounted future reward. As the discount factor approached 1 (i.e. no temporal discounting), this model converged to long-term reward maximization, equivalent to MVT. Additional parameters were added to the model to test four specific hypotheses for suboptimal foraging behavior: i) subjective costs associated with leaving a patch, in which the value of leaving was reduced by a “cost” term; ii) diminishing marginal returns, in which the subjective utility of a reward increased sublinearly with respect to the reward magnitude; iii) biased time perception, which assumes that animals underestimate post-reward delays or overestimate pre-reward delays (Bateson and Kacelnik, 1996; Blanchard et al., 2013; Stephens and Anderson, 2001); and iv) temporal discounting. For each model, group level parameters and parameters for each individual rat were fit simultaneously using an expectation-maximization algorithm (Huys et al., 2011). Parameters were fit to each experiment separately (one set of parameters for both conditions in each experiment). Model predictions were calculated separately for each rat, using the rat’s individual parameters. Full details for all models, fitting procedures, and model comparison can be found in the methods.

Subjective costs to leave a patch and diminishing marginal returns for larger rewards have explained suboptimal foraging behavior in prior studies that have manipulated opportunity costs (e.g. travel time or pre-reward delays) and depletion rate (Carter and Redish, 2016; Constantino and Daw, 2015; Wikenheiser et al., 2013). However, these factors are insensitive to the placement of time delays (pre- vs. post-reward) and thus, cannot explain the preference for more immediate rewards. Consistent with prior studies, these hypotheses explained overharvesting in the travel time and Depletion Rate Experiments (Fig. S3A-B), but they failed to explain time preferences in the pre-vs-post foraging experiment (Fig. S3D).

We next examined whether biased time perception and temporal discounting could explain suboptimal foraging behavior across all tasks. Three implementations of each hypothesis were tested: for biased time perception, linear underestimation of post-reward delays (*postDelay* = *α* * *postDelay)*, non-linear underestimation of post-reward delays 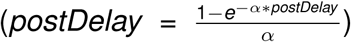, and overestimation of pre-reward delays (*preDelay* = *α* * *preDelay)*; for temporal discounting, exponential (*value* = *e*^*-β*time*^ * *reward)*, standard hyperbolic (*value* = *reward/*(1 + *k* * *time*)), and quasi-hyperbolic — a more flexible form of hyperbolic discounting, formalized as two competing exponential discounting systems (*value* = [*ω* \**e*^*-β*time*^ + (1 − *ω*) * *e*^*-δ*time*^] \**reward*; Laibson 1997; McClure et al. 2007). All models tested explained rat behavior in a subset of experiments, but only one model, quasi-hyperbolic discounting, explained rat behavior across all tasks (Figs. 3A-E, S4-S5).

**Figure 3.**
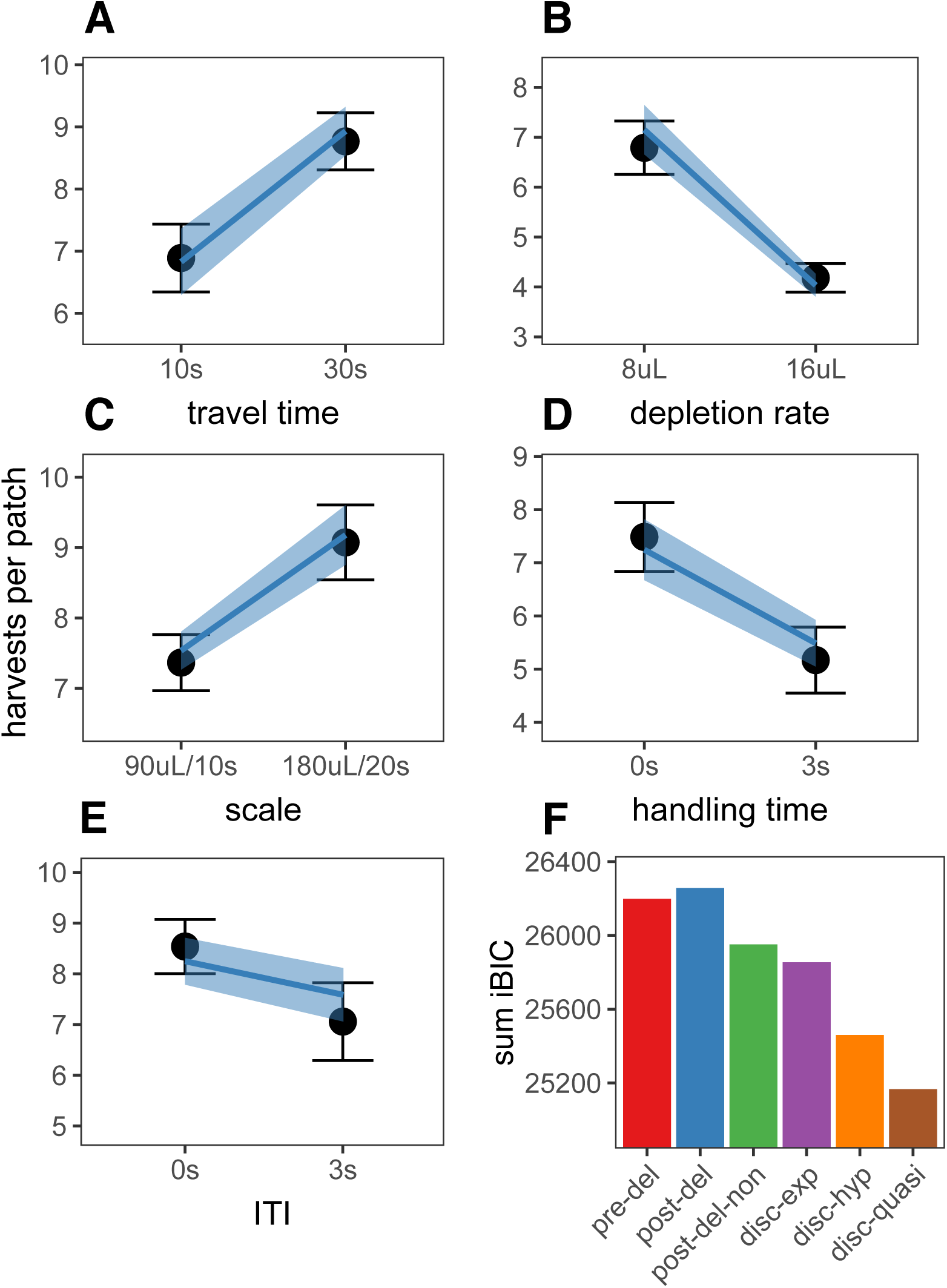
A-E) Predictions of the best fit quasi-hyperbolic discounting model to all foraging tasks. Points and error bars represent mean ± standard deviation of the means for each individual rat; lines and ribbon represent the mean ± standard deviation of the means of the model-predicted behavior for each individual rat. F) The sum of iBIC scores across all foraging tasks for each model. Pre-del = linear overestimation of pre-reward delays, post-del = linear underestimation of post-reward delays, post-del-non = nonlinear underestimation of post-reward delays, disc-exp = exponential discounting, disc-hyp = hyperbolic discounting, disc-quasi = quasi-hyperbolic discounting.

To determine which model provided the best quantitative fit, we compared the group-level Bayes Information Criterion (integrated BIC or iBIC; Huys et al. 2011, 2012) of all models in each of the foraging tasks. The quasi-hyperbolic discounting model had the lowest cumulative score across all foraging tasks (sum of iBIC across tasks; Fig. 3F) and also had the lowest iBIC score in all individual foraging experiments except for the Depletion Rate Experiment, in which it ranked second to the hyperbolic discounting model (iBIC_hyperbolic_ = 5726.858, iBIC_quasi-hyperbolic_ = 5732.990; Fig. S6). Even in this experiment, the negative log marginal likelihood (-LML; group level negative log likelihood) of the quasi-hyperbolic discounting model was lower than that of the hyperbolic discounting model (-LML_hyperbolic_ = 5710.579, -LML_quasi_ = 5700.429), indicating that the quasi-hyperbolic model provided a better absolute fit, but iBIC placed a greater penalty on the quasi-hyperbolic discounting model due to additional parameters (4 for quasi-hyperbolic vs. 2 for hyperbolic).

Next, we tested whether the model that provided the best fit to foraging behavior, quasi-hyperbolic discounting, could also explain behavior in the intertemporal choice task. For this task, each series of 10 free choices was modeled as a separate episode, such that the value of the first choice was equal to discounted reward across all 10 choices, the value of the second choice equal to discounted reward across the remaining 9 choices, and so on (see abbreviated state space diagram in Fig. S2B). The episode ended at the 10th choice, and did not consider rewards in future games. We tested all biased time perception and temporal discounting models in this task. All three temporal discounting models had lower iBIC scores than all three biased time perception models. Again, the quasi-hyperbolic discounting model had the lowest iBIC score among all candidates (iBIC_exp_ = 89712.79, iBIC_hyp_ = 89712.35, iBIC_quasi_ = 89679.39; Fig. 4B, quasi-hyperbolic model predictions in Fig. 4A).

**Figure 4.**
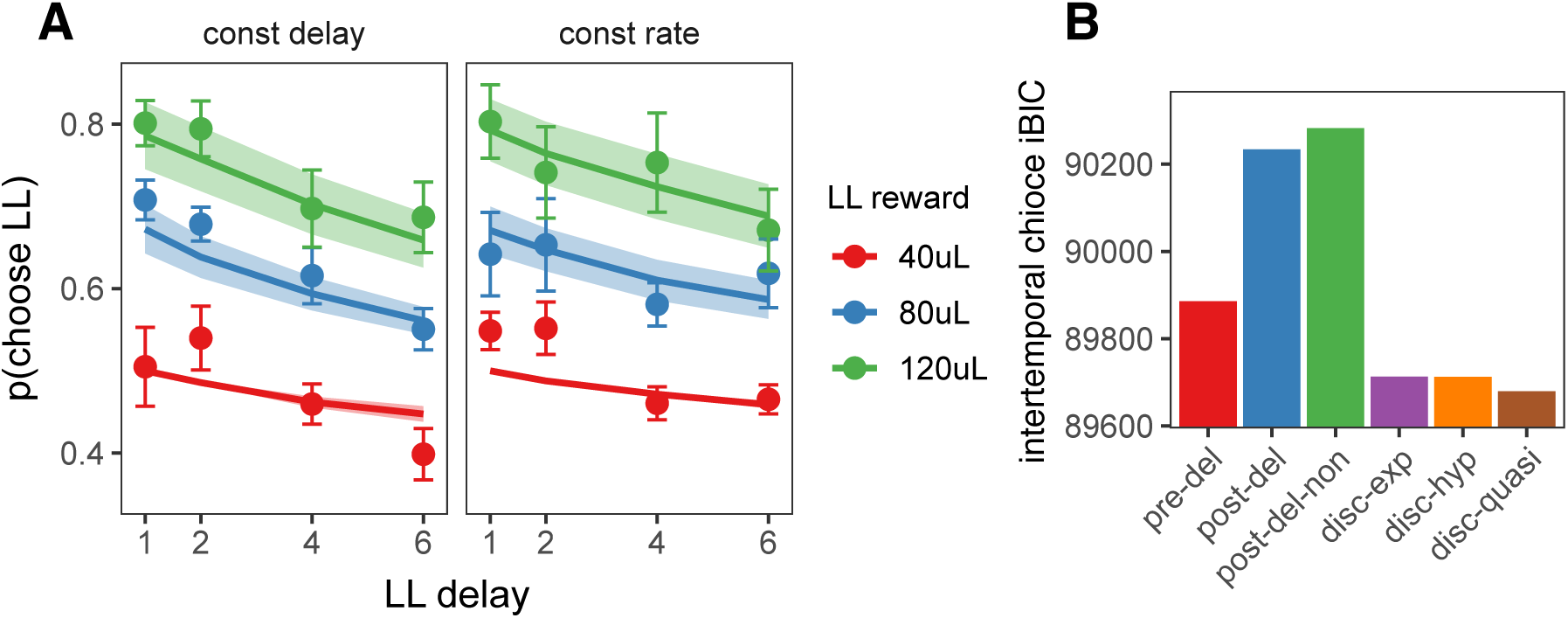
A) Quasi-hyperbolic model predictions for the intertemporal choice task. Points and error bars represent the mean ± standard error of individual rat behavior; lines and ribbon represent mean ± standard error of model predicted behavior for each individual rat. B) The iBIC score for each model for the delay discounting experiment. Pre-del = linear overestimation of pre-reward delays, post-del = linear underestimation of post-reward delays, post-del-non = nonlinear underestimation of post-reward delays, disc-exp = exponential discounting, disc-hyp = hyperbolic discounting, disc-quasi = quasi-hyperbolic discounting.

The quasi-hyperbolic discounting model performed the best among all candidate models in both tasks. If quasi-hyperbolic discounting reflects a common explanation for suboptimal decision-making, then it might be expected that animals should exhibit similar discount functions across tasks. To test the external validity of the quasi-hyperbolic discounting model, we examined how well the discount function that best fit foraging behavior of each individual rat could predict its intertemporal choice behavior, and vice-versa. For this analysis, data from each task were separated into three subsets. The quasi-hyperbolic discounting model was fit to two subsets of data from one task, then the negative log likelihood (-LL) of the data was assessed on the left out sample from both tasks. This process was repeated such that each subset served as the left out sample. To determine which discount function provided the better fit to data from each task, we calculated the difference in -LL of the left out sample between the model fit to intertemporal choice data and the model fit to foraging data (-LL difference = -LL_itc_ - -LL_forage_). Since smaller -LL indicates a better fit, a positive -LL difference indicates that the discount function fit to foraging data provided a better fit (i.e. the foraging -LL was lower than the intertemporal choice -LL). For the foraging task, discounting functions fit to foraging data provided a better fit than discounting functions fit to intertemporal choice data for 7/8 rats. Interestingly, for the intertemporal choice task, discounting functions fit to foraging data provided a better fit than discounting functions fit to intertemporal choice data for the majority of rats (5/8; Fig. 5C). Taken together, these findings provide support for the idea that foraging and intertemporal choice can be described by a common discount function.

**Figure 5.**
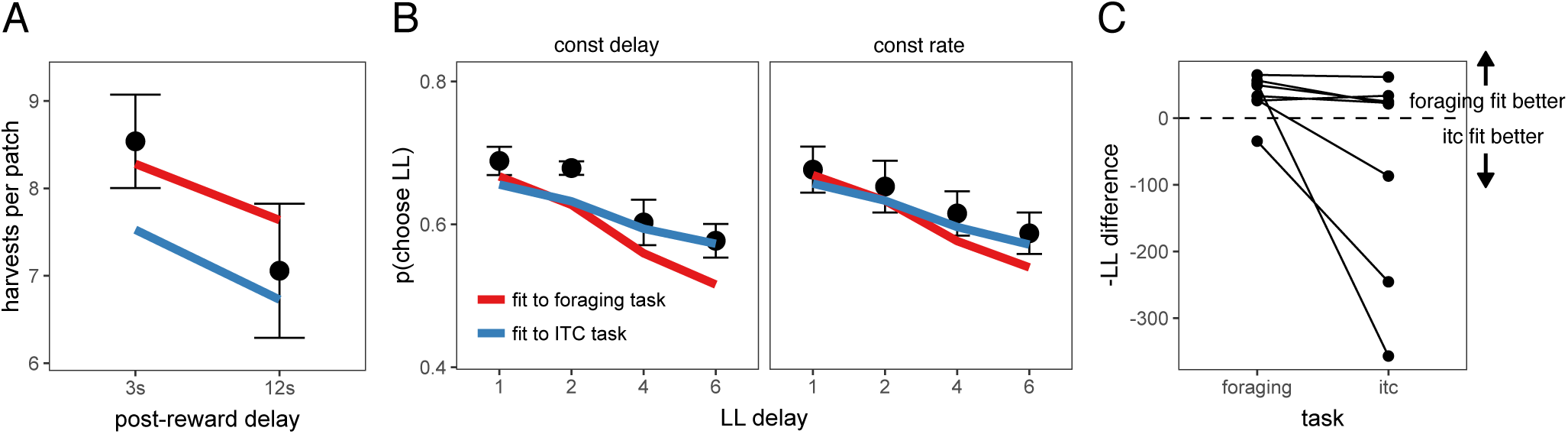
A) Predicted foraging behavior for quasi-hyperbolic model parameters fit to either the foraging task (red line) or delay discounting task (DD; blue line). Black points and error bars represent mean ± standard error of rat data. B) Predicted intertemporal choice behavior for quasi-hyperbolic model parameters fit to data from either the foraging or delay discounting task, plotted against rat behavior. C) The difference in negative log likelihood of the left out sample of foraging data (left) or intertemporal choice data (right) between parameters fit to the intertemporal choice task and parameters fit to the foraging task. A negative -LL difference indicates the negative log likelihood of the data for parameters fit to the intertemporal choice task was lower than for parameters fit to the foraging task. Each point and line represents data from individual rats.

## Discussion

In foraging studies, animals exhibit behavior that conforms qualitatively to predictions made by optimal foraging theory (i.e., the MVT), choosing to leave a patch when its value falls below that of the average expected value of other(s) available in the environment. However, an almost ubiquitous finding is that they overharvest, leaving a patch when its value falls to a value lower than the one predicted by MVT. Given that the rewards available within the current patch are generally available sooner than those at other patches due to travel time, one interpretation of overharvesting is that this reflects a similarly prevalent bias observed in intertemporal choice tasks, in which animals consistently show a greater preference for smaller more immediate rewards over later delayed rewards than would be predicted by optimal (i.e,. exponential) discounting of future values. However, in prior studies, models of intertemporal choice behavior have been poor predictors of foraging behavior (Blanchard and Hayden, 2015; Carter and Redish, 2016). Here, we show that in a carefully designed series of experiments, rats exhibit similar time preferences in foraging and intertemporal choice tasks, and that a quasi-hyperbolic discounting model can explain the rich pattern of behaviors observed in both tasks.

The foraging behavior we observed was consistent with previous studies of foraging behavior in rats, monkeys, and humans, while also revealing novel aspects of overharvesting behavior. Consistent with prior studies, rats stayed longer in patches that yielded greater rewards, stayed longer in all patch types when the cost of traveling to a new patch was greater, left patches earlier when rewards depleted more quickly, and consistently overharvested (Constantino and Daw, 2015; Hayden et al., 2011; Kane et al., 2017). Our experiments also demonstrated that in certain environments rats violate qualitative predictions of MVT. Rats overharvested more when reward amount and delay were increased, even though reward rate was held constant, and they were differentially sensitive to whether the delay was before the receipt of the proximal reward or following its delivery. These findings supported the conjecture that overharvesting is related to time preferences. When we directly tested rats sensitivity to post-reward delays, we found that contrary to previous studies addressing time preferences (Bateson and Kacelnik, 1996; Blanchard et al., 2013; Mazur, 1991; Stephens and Anderson, 2001), rats were sensitive to post-reward delays in both the foraging and the intertemporal choice tasks.

The idea that animals exhibit similar decision biases in foraging and intertemporal choice paradigms, and that these biases can be explained by a common model of discounting, is in conflict with prior studies that found that animals are better at maximizing long-term reward rate in foraging than in intertemporal choice tasks, and that delay discounting models of intertemporal choice tasks are poor predictors of foraging behavior (Blanchard and Hayden, 2015; Carter et al., 2015; Carter and Redish, 2016; Stephens, 2008). It has been argued that animals may perform better in foraging tasks because decision-making systems have evolved to solve foraging problems rather than two-alternative intertemporal choice problems (Blanchard and Hayden, 2015; Stephens, 2008; Stephens et al., 2004). However, results from the present study contradict the notion that animals use different strategies in these different task structures. There are two potential explanations for why temporal discounting models have failed to predict foraging behavior in prior studies: i) prior studies have only tested single-parameter exponential and hyperbolic discounting functions, whereas the present study also tested the more flexible quasi-hyperbolic discounting function; and ii) in most of these studies, models of intertemporal choice tasks have only considered the most proximal reward (the reward received as a consequence of the decision at hand). This assumption seems appropriate as, In most intertemporal choice tasks, opportunities for future rewards do not depend on the current decision, so the value of rewards received for future decisions are equal for both the SS and LL rewards. However, adding the value of future rewards increases the perceived value of both options, which may change the estimate of the discount factors. Since, in foraging tasks, future opportunities for reward depend on current decisions, it is critical for foraging models to include all future rewards into estimates of reward value. For this reason, comparing discount functions fit to intertemporal choice models that consider all future reward may provide better estimates of foraging behavior than discount function fit to intertemporal choice models that only consider rewards from the most proximal decision.

Although quasi-hyperbolic discounting provided the best singular explanation for rat behavior across tasks, many of the models tested were capable of explaining some of the biases exhibited by rats. Thus, we cannot exclude the possibility that subjective costs, diminishing marginal utility, and/or biased estimation of time intervals may independently contribute to suboptimal decision-making. Importantly, our data indicate that quasi-hyperbolic discounting may provide a link between foraging and intertemporal choice tasks, and it highlights the importance of future work considering the source of time preferences. These observations are buttressed by recent theoretical work demonstrating that the appearance of time preferences in intertemporal choice tasks can emerge rationally from a value construction process by which estimates increase in variability with the delay until reward receipt — an account that shares features with the short-term rate maximization hypotheses (Stephens et al., 2004). Under this account, “as-if” discounting is hyperbolic when variability increases linearly with delay (Gabaix and Laibson, 2017). Further, a sequential sampling model of two-alternative forced choice (Bogacz et al., 2006), parameterized such that outcome delay scales variability in this way, has recently been shown to capture key dynamical features of both patch foraging (Davidson and El-Hady, 2018) and hyperbolic discounting in intertemporal choice (Hunter et al., 2018). Future work should build on these findings to explore directly whether the common biases identified here reflect a core computation underlying sampling and decision-making under uncertainty and across time.

## Methods

### Animals

Adult Long-Evans rats were used (Charles River, Kingston, NY). One group of eight rats participated in the scale, travel time, depletion rate, and handling time experiments (in that order), a different set of eight rats were tested on the post-reward delay foraging experiment then the delay discounting task. Rats were housed on a reverse 12 h/12 h light/dark cycle. All behavioral testing was conducted during the dark period. Rats were food restricted to maintain a weight of 85-90% ad-lib feeding weight, and were given ad-lib access to water. All procedures were approved by the Princeton University and Rutgers University Institutional Animal Care and Use Committee.

### Foraging Task

Animals were trained and tested as in Kane et al. (2017). Rats were first trained to lever press for 10% sucrose water on an FR1 reinforcement schedule. Once exhibiting 100+ lever presses in a one hour session, rats were trained on a sudden patch depletion paradigm — the lever stopped yielding reward after 4-12 lever presses — and rats learned to nose poke to reset the lever. Next rats were tested on the full foraging task.

A diagram of the foraging task is in Fig. S1. On a series of trials, rats had to repeatedly decide to lever press to harvest reward from the patch or to nose poke to travel to a new, full patch, incurring the cost of a time delay. At the start of each trial, a cue light above the lever and inside the nose poke turned on, indicating rats could now make a decision. The time from cues turning on until rats pressed a lever or nose poked was recorded as the decision time (DT). A decision to harvest from the patch (lever press) yielded reward after a short pre-reward delay (referred to as the handling time delay, simulating the time to “handle” prey after deciding to harvest). Reward (sucrose water) was delivered when the rat entered the reward magazine. The next trial began after an inter-trial interval (ITI). To control the reward rate within the patch, the length of the ITI was adjusted based on the DT of the current trial, such that the length of all harvest trials was equivalent. With each consecutive harvest, the rat received a smaller volume of reward to simulate depletion from the patch. A nose poke to leave the patch caused the lever to retract for a delay period simulating the time to travel to a new patch. After the delay, the opposite lever extended, and rats could harvest from a new, replenished patch.

Details of the foraging environment for each experiment can be found in Table 1. For each experiment, rats were trained on a specific condition for 5 days, then tested for 5 days. Conditions within experiments were counterbalanced. Rat foraging behavior was assessed using mixed effects models. In the Travel Time Experiment, we assessed the effect of starting volume of the patch and the travel time on number of harvests per patch, with random intercepts and random slopes for both variables across subjects. In all other foraging experiments, we assessed the effect of experimental condition on harvests per patch, with random intercepts and random effect of experimental condition across subjects.

### Two-alternative choice task

Rats were immediately transferred from the foraging task to the two alternative choice task with no special training; rats were given three 2-hour sessions to learn the structure of the new task. This task consisted of a series of episodes that lasted 20 trials. At the beginning of each episode one lever was randomly selected as the shorter-sooner lever, yielding 40 µL of reward following a 1 s delay. The other lever (larger-later lever) was initialized to yield a reward of 40, 80, or 120 µL after a 1, 2, 4 or 6 s delay. For the first 10 trials of each episode, only one lever extended, and rats were forced to press that lever to learn its associated reward value and delay. The last four forced trials (trials 7-10) were counterbalanced to reduce the possibility of rats developing a perseveration bias. For the remaining 10 trials of each episode, both levers extended, and rats were free to choose the option they prefer. At the beginning of each trial, cue lights turned on above the lever indicating rats could now make a decision. Once the rat pressed the lever, the cue light turned off, and the delay period was initiated. A cue light turned on in the reward magazine at the end of the delay period, and rats received reward as soon as they entered the reward magazine. Reward magnitude was cued by light and tone. Following reward delivery, there was an ITI before the start of the next trial. At the completion of the episode, the levers retracted, and rats had to nose poke to begin the next episode, which reset the larger-later reward and delay.

Two-alternative choice data was analyzed using a mixed effects logistic regression, examining the the effect of larger-later reward value, larger-later delay, and task condition on rats choices, with random intercepts and random effects for all three variables. Custom contrasts were tested using the phia package in R (de Rosario-Martinez, 2015), using Holm’s method to correct for multiple comparisons.

### Foraging Models

All models were constructed as continuous time semi-markov processes. This provided a convenient way to capture the dynamics of timing in both tasks, such as slow delivery and consumption of reward (up to 6 s for the largest rewards). To model the foraging task, each event within the task (e.g. cues turning on/off, lever press, reward delivery, etc.) marked a state transition (abbreviated state space diagram in Fig. S2). All state transitions were deterministic, except for decisions to stay in vs. leave the patch, which occurred in ‘decision‘ states (the time between cues turning on at the start of the trial and rats performing a lever press or nosepoke). In decision states, a decision to stay in the patch transitioned to the handling time state, then reward state, ITI state, and to the decision state on the next trial. A decision to leave transitioned to the travel time state, then to the first decision state in the patch. Using the notation of Bradtke and Duff (1995), the value of staying in state *s, Q*(*stay, s*), is the reward provided for staying in state *s, R*(*stay, s*), plus the discounted value of the next state:

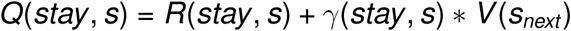

where *γ*(*stay, s*) is the discount applied to the value of the next state for staying in state *s*, and *V* (*s*_*next*_) is the value of the next state in the patch. For all non-decision states, rats did not have the option to leave the patch, so for these states, *V* (*s*) = *Q*(*stay, s*). For decision states, the value of the state was the greater of *Q*(*stay, s*) and *Q*(*leave*).

For simplicity, we assume the time spent in a given state is constant, calculated as the average amount of time a given rat spent in the state. Under this assumption, *R*(*stay, s*) is the reward rate provided over the course of the state, *r* (*s*), multiplied by the time spent in that state *T* (*s*), discounted according to discount factor *β*:

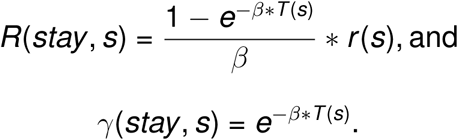

The value of leaving a patch, *Q*(*leave*), was equal to the discounted value of the first state in the next patch, *V* (*s*_*first*_):

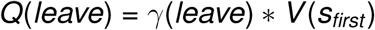

where *γ*(*leave*) is the discount factor applied to the next state in the first patch. Assuming no variance in the travel time *τ, γ*(*leave*) = *e*^*-β*τ*^. Per MVT, we assumed rats left patches at the first state in the patch in which *Q*(*stay, s*) ≤ *Q*(*leave*). To model variability in the trial at which rats left patches, we added gaussian noise to *Q*(*leave*). As decisions within each patch are not independent, the patch leaving threshold did not vary trial-by-trial, but rather patch by patch, such that the cumulative probability that a rat has left the patch by state *s, π*(*leave, s*), was the probability that *Q*(*stay, s*) ≤ *Q*(*leave*) + ϵ, where ϵ ∼ 𝒩 (0, *σ*^2^), with free parameter *σ*.

The optimal policy for a given set of parameters was found using dynamic programming. Optimal foraging behavior is to maximize undiscounted long-term reward rate. Optimal behavior was determined by fixing the discount rate factor *β* = .001 and assuming no decision noise (ϵ = 0). Optimal behavior was determined for each rat, in which the time spent in each state was taken from a given rat’s data. For each model, we fit both group level parameters and individual parameters for each rat using an expectation-maximization algorithm (Huys et al., 2011).

To model subjective costs, a free parameter *c* representing an aversion to leaving the patch was subtracted from the leaving threshold (Carter and Redish, 2016; Wikenheiser et al., 2013):

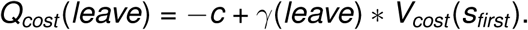

To investigate whether diminishing marginal returns could explain rats overharvesting behavior, we tested models in which the utility of a reward received in the task increased in a sublinear fashion with respect to the magnitude of the reward. Two different utility functions were tested: a power law function and a steeper constant relative risk aversion (CRRA) utility function that became increasingly risk averse with larger rewards, both with free parameter *η*:

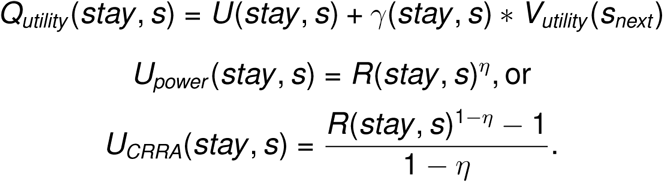

To examine linear and nonlinear underestimation of post-reward delays, respectively, the time spent in post-reward delay (ITI) states was transformed, with free parameter *α*:

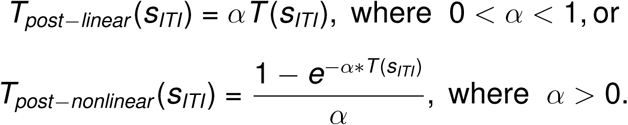

Similarly, for overestimation of pre-reward delays, the handling time and travel time were transformed:

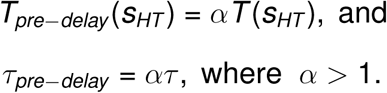

For the exponential discounting model, *β* was fit as a free parameter. As standard hyperbolic discounting cannot conveniently be expressed recursively, this model was implemented using the µAgents model described by Kurth-Nelson and Redish (2009). The value functions of the overall model, 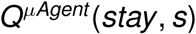 and 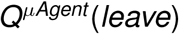, were the average of 100 µAgents, each with their own exponential discount factor *β*_*i*_, and thus individual reward functions *R*_*i*_(*stay, s*), discount functions *γ*_*i*_(*stay, s*) and *γ*_*i*_(*leave*), and value functions *Q*_*i*_(*stay, s*), *Q*_*i*_(*leave*), and *V*_*i*_(*s*):

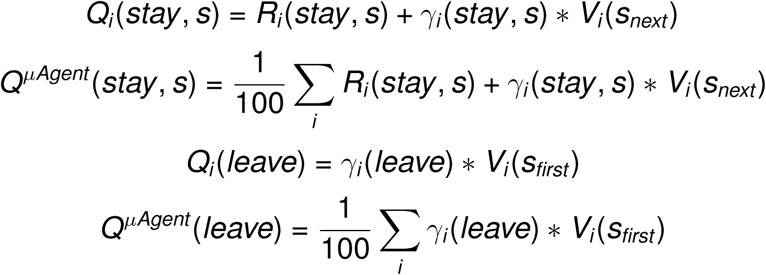

If the µAgent discount factors, *β*_*i*_, were drawn from an exponential distribution with rate parameter *λ* > 0, the discounting function of the overall model approximated the standard hyperbolic discount function, *reward/*(1+*k delay)*, with discount rate *k* = 1*/λ*. This relationship is presented in Fig. S8. *λ* was fit as a free parameter.

Quasi-hyperbolic discounting was originally formulated for discrete time applications (Laibson, 1997). We used the continuous time formulation from McClure et al. (2007), in which the value functions of the overall model were the weighted sum of two exponential discount systems, a steep discounting *β* system that prefers immediate rewards and a slower discounting *δ* system, each with their own reward functions, *R*_*β*_(*stay, s*) and *R*_*δ*_(*stay, s*), and discount functions *γ*_*β*_(*stay, s*), *γ*_*β*_(*leave*), *γ*_*δ*_(*stay, s*), and *γ*_*δ*_(*leave*):

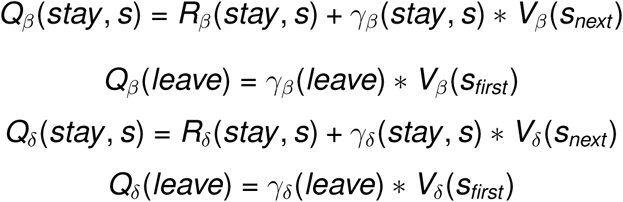

The value function of the overall quasi-hyperbolic discounting model were:

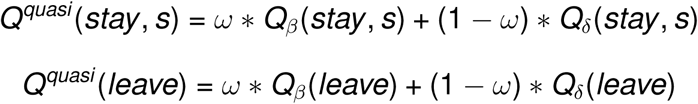

where 0 < *ω* < 1 was the weight of the *β* system relative to the *δ* system. *β, δ*, and *ω* were all free parameters.

### Intertemporal Choice Task Models

Similar to the foraging task, events within the intertemporal choice task marked state transitions, and all state transitions were deterministic except for decisions to choose the smaller-sooner option (SS) or larger-later option (LL), which occurred only in decision states (abbreviated state space diagram in Fig S2B). From decision states, animals transitioned to delay, reward, and post-reward delay (ITI) states for the chosen option — the delay, reward and ITI for the SS and LL options were represented by separate states. The value of choosing SS or LL in decision state *s* is the discounted value of the next state, the following delay state:

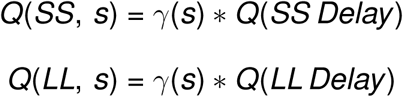

The value of delay states were the discounted value of the reward state for that action, the value of reward states were the reward for that action plus the discounted value of the ITI state for that action, and the value of ITI states were the discounted value of the next decision state:

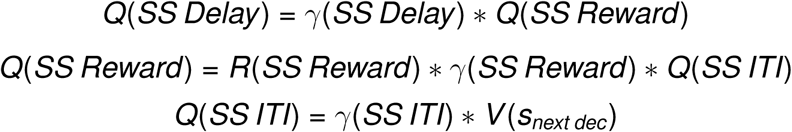

where the value of the next decision state, *V*(*s*_*next dec*_) is the greater of *Q*(*SS, s*_*next dec*_) and *Q*(*LL, s*_*next dec*_). Decisions were made using a softmax rule, with the probability of choosing the LL option in decision state *s* defined as:

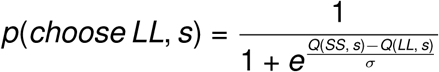

with temperature parameter *σ*, a free parameter that determines decision noise. The underestimating post-reward delays and temporal discounting models were implemented as they were in the foraging task.

### Model Comparison

All models had two parameters except for the quasi-hyperbolic discounting model, with four. To determine the model that provided the best fit to the data, while accounting for the increased flexibility of the quasi-hyperbolic discounting model, we calculated the Bayes Information Criterion over the group level parameters (iBIC) (Huys et al., 2011; MacKay, 2003). iBIC penalizes the log marginal likelihood, *logp*(*D* | *θ*), which is the integral of the log likelihood of the data *D* over the distribution of group level parameters *θ*, for model complexity. Complexity is determined by the number of parameters *k,* and the size of the penalty depends on the total number of observations, *n*:

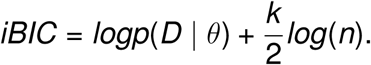

As in Huys et al. (2011), we use a Laplace approximation to the log marginal likelihood:

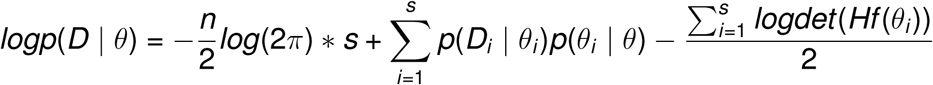

where *s* is the number of subjects, and *Hf* (*θ*_*i*_) is the hessian matrix of the likelihood for subject *I* at the individual parameters *θ*_*i*_.

To compare the fit of the quasi-hyperbolic discounting model across the foraging and intertemporal choice tasks, data from each task was separated into thirds. The quasi-hyperbolic discounting model was fit to 2 of the samples from each task using maximum likelihood estimation (fitting only individual parameters for each rat). The log likelihood of the data from the left out sample was evaluated. This process was repeated three times, leaving out each of the samples once, and we took the sum of the likelihood of the three left out samples. As the structure of variability was different between the foraging model (variability in the patch leaving threshold) and intertemporal choice models (softmax decision noise), to compare the discount function fit to the foraging task on intertemporal choice data, a new noise parameter was fit to the intertemporal choice data (and vice-versa). We report the difference in the log likelihood of the data using parameters fit to the intertemporal choice task and of the log likelihood using parameters fit to the foraging task (Fig. 5.

## Acknowledgements

We thank Dr. Gary Aston-Jones for helpful discussions. This work was supported by NIH grant F31MH109286 (GAK) and the Princeton Program in Cognitive Science.

## Supplemental Figures

**Figure S1.**
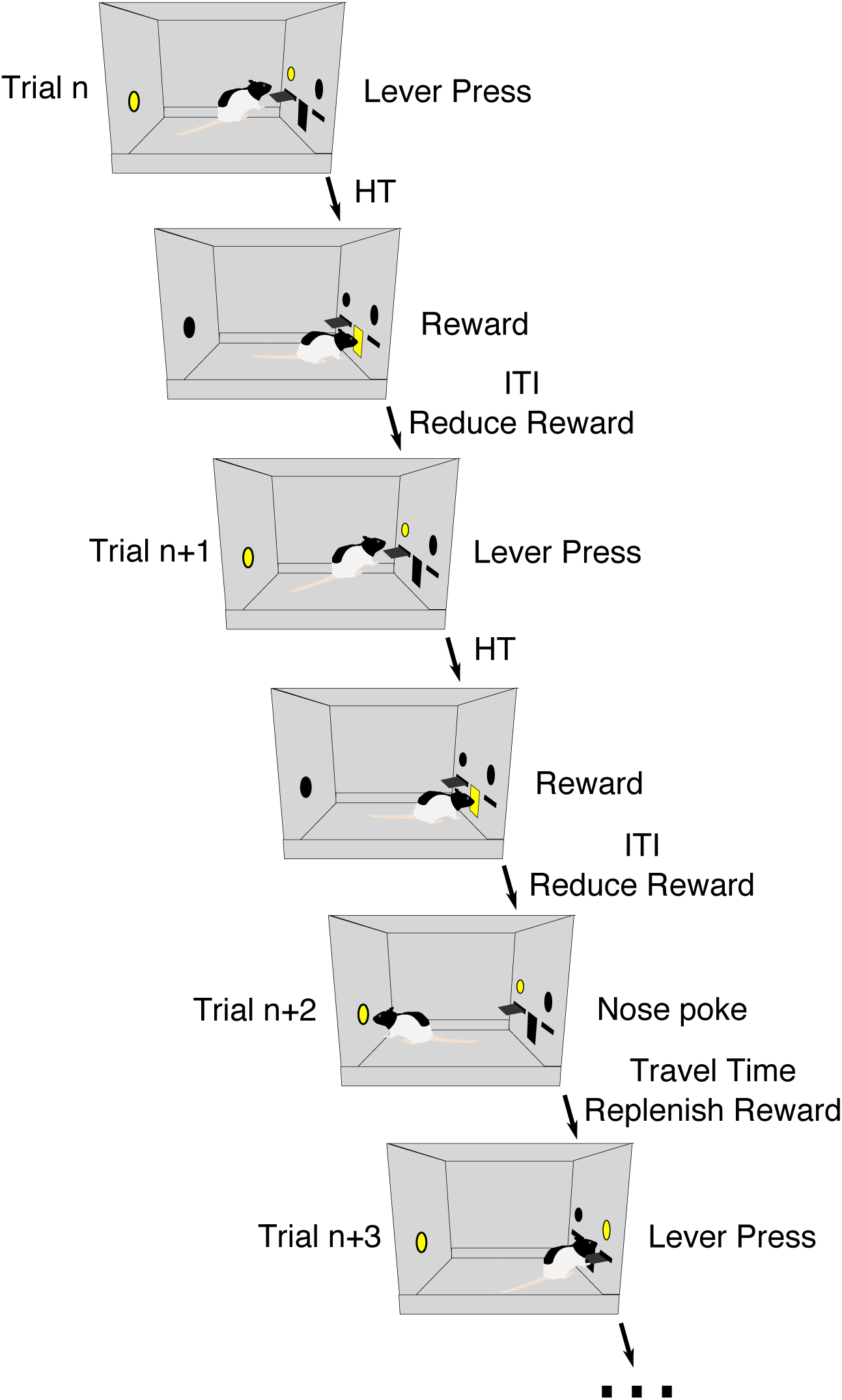
Diagram of the foraging task. Rats press a lever to harvest reward from the patch then receive reward in an adjacent port following a handling time delay. After receiving reward, there is an inter-trial interval (post-reward delay) before rats can make their next decision. Rats can leave the patch by nose poking in the back of the chamber (trial n+2), which initiates a delay simulating time to travel to the next patch, after which, rats can harvest from a new replenished patch.

**Figure S2.**
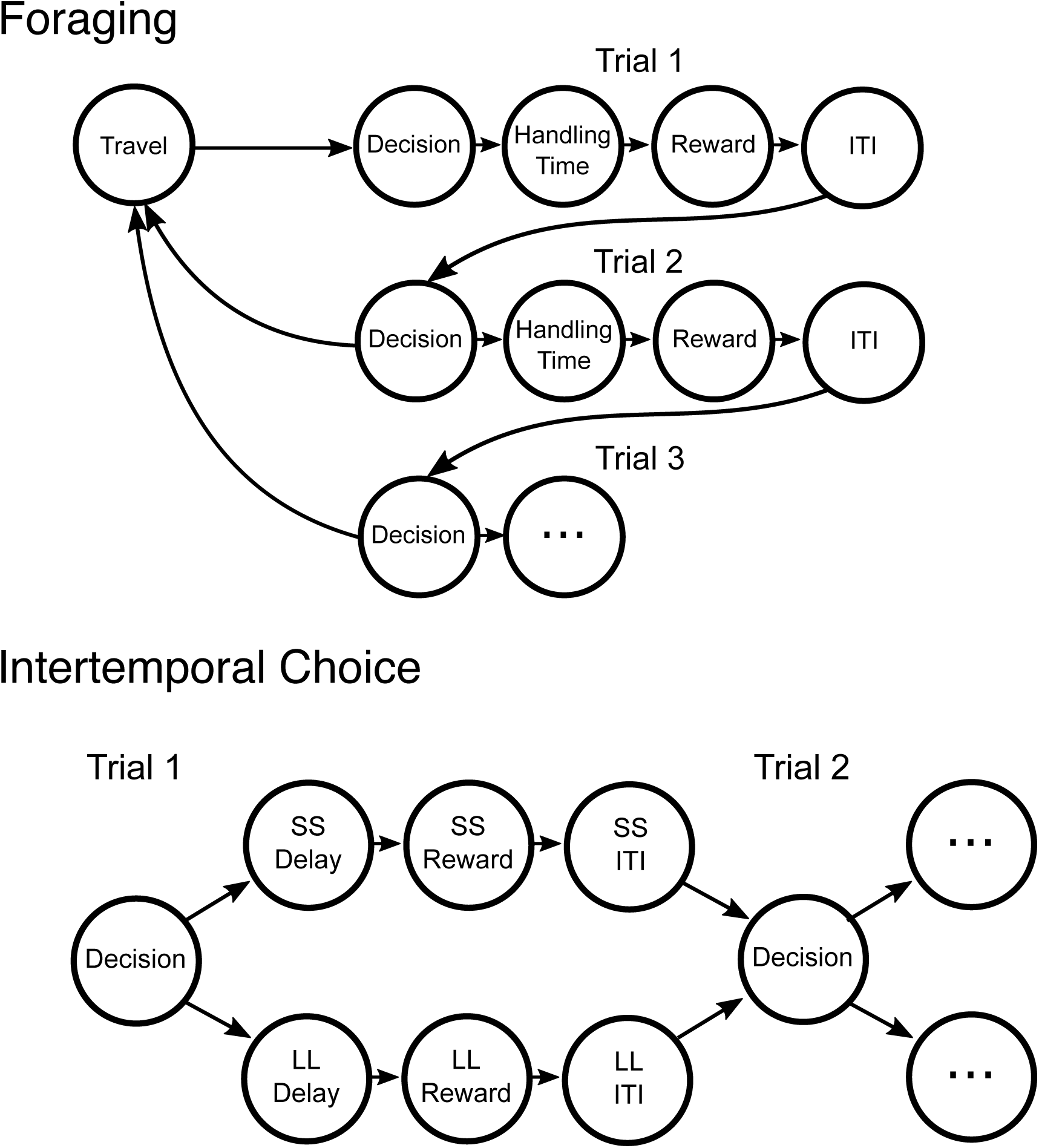
State space diagram for the semi-markov model of the foraging (top) and intertemporal choice (bottom) tasks. In the foraging task, decisions to stay vs. leave are made in Decision states. A Decision to stay causes a transition to the handling time, then reward, ITI, and to the Decision state on the next trial. Reward is delivered uniformly throughout time spent in the each reward state. Reward depletion is achieved via shorter time spent in the reward state (resulting in longer stay in the ITI state). A Decision to leave causes a transition to the travel state, then to the first trial of the patch. In the intertemporal choice task, decisions made in Decision states cause transition to the Delay, Reward, and ITI states for the option chosen (either SS or LL), then back to the next Decision state. The model consisted of 10 consecutive trials— the number of free choice trials.

**Figure S3.**
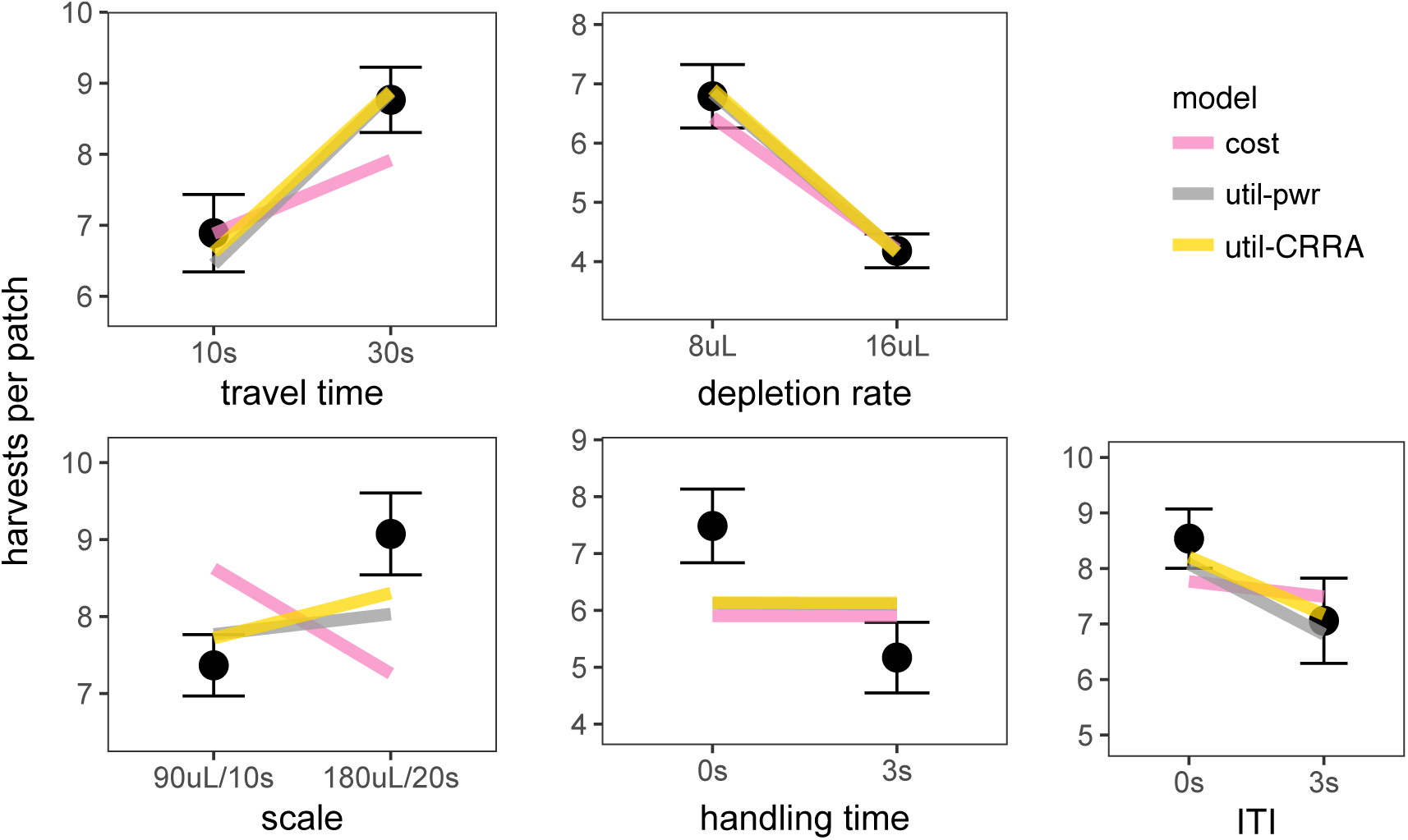
Predictions of the best fit subjective cost and diminishing marginal returns models (power law = util-pwr; constant relative risk aversion = util-CRRA). Black points and error bars represent mean ± standard error of observed behavior. Colored lines represent the mean model predicted behavior across rats.

**Figure S4.**
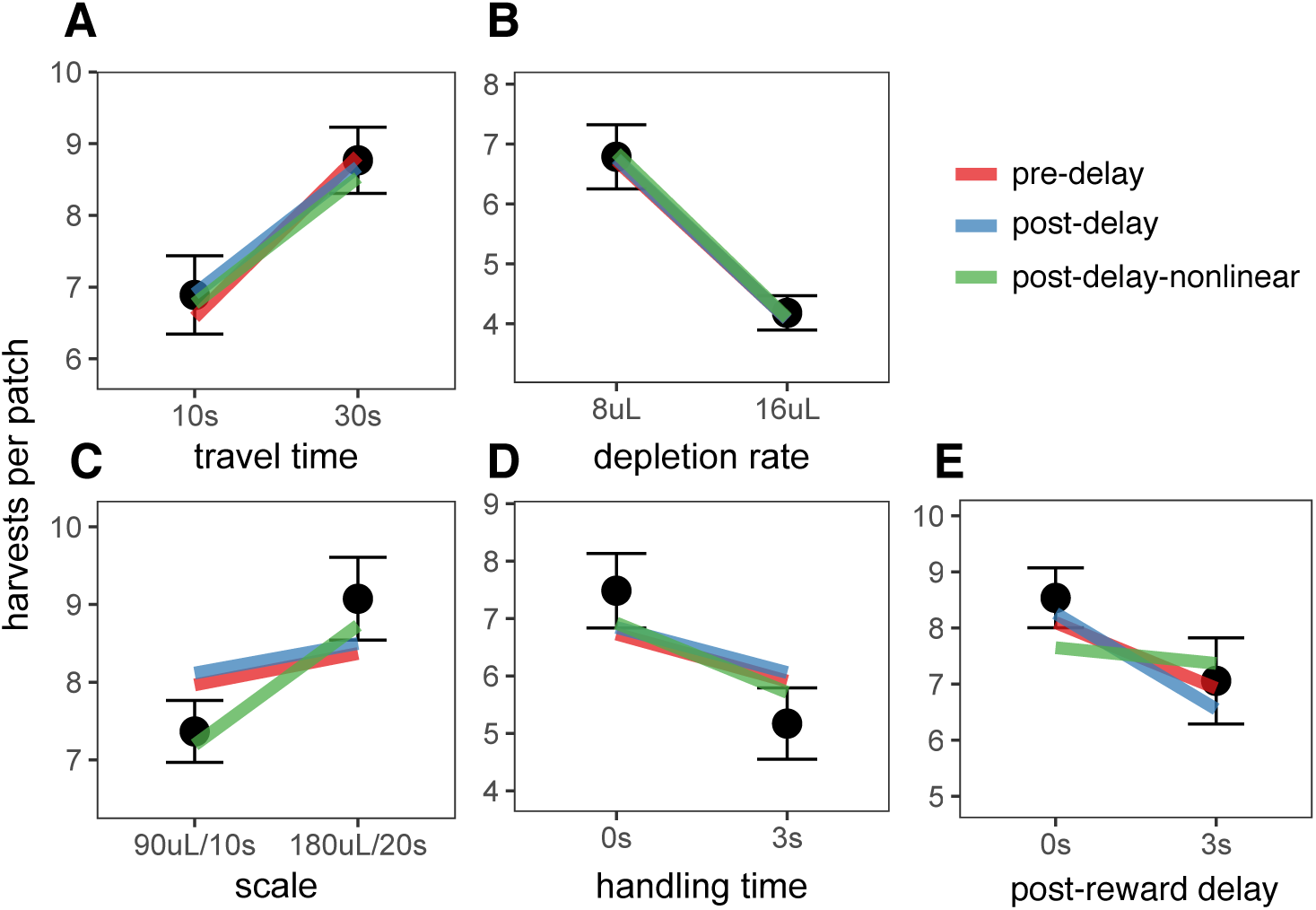
Predictions of the best fit models of overestimation of pre-reward delays (pre-delay), linear underestimation of post-reward delays (post-delay), and nonlinear underestimation of post-reward delays (post-delay-nonlinear). Points and errorbars are the mean ± standard deviation of rat behavior, colored lines represent the mean model predicted number of harvests across all rats.

**Figure S5.**
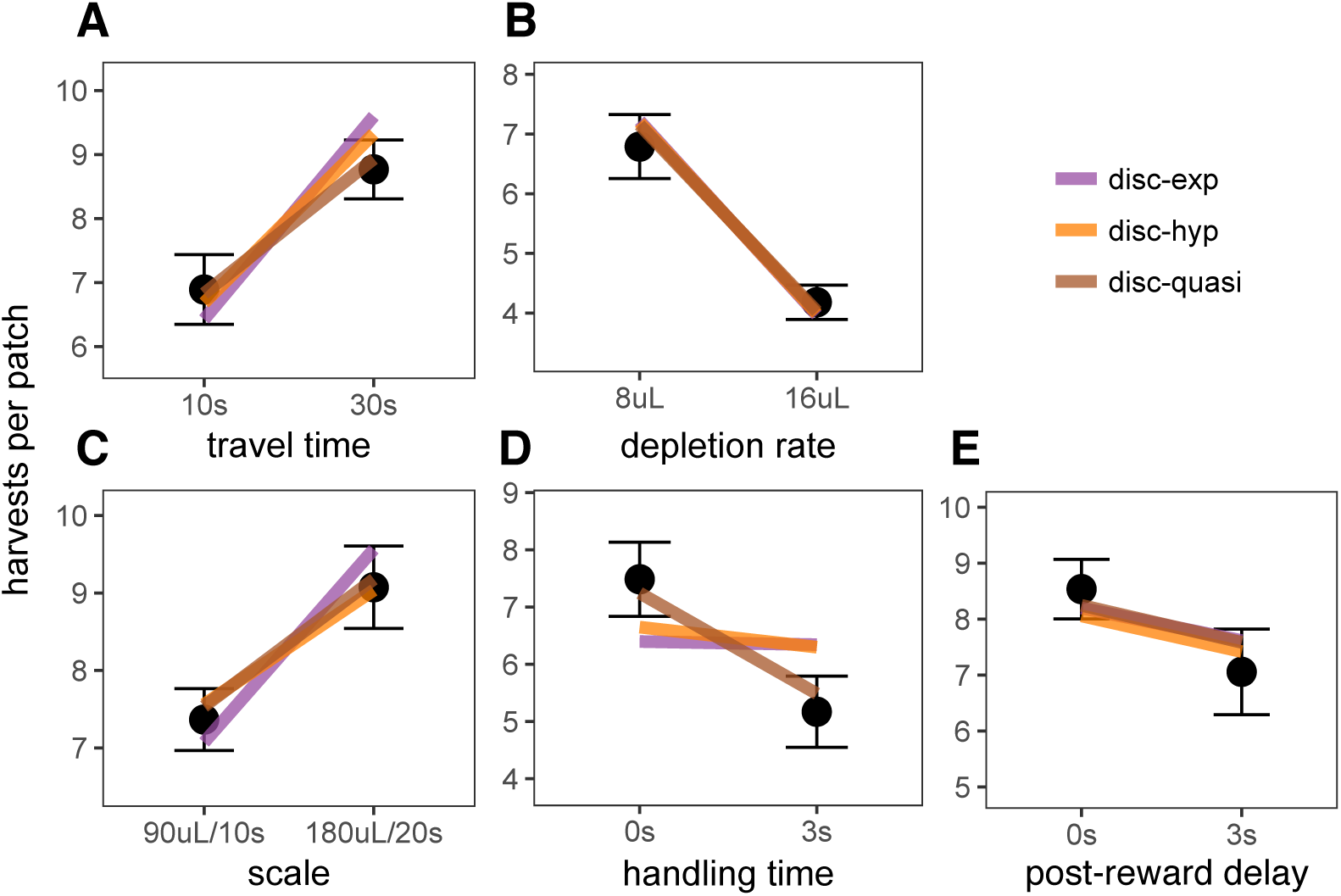
Predictions of the best fit exponential discounting model (disc-exp), hyperbolic discounting model (disc-hyp), and quasi-hyperbolic discounting model (disc-quasi). Points and error bars are the mean ± standard deviation of rat behavior; colored lines represent the mean predicted number of harvests across all rats.

**Figure S6.**
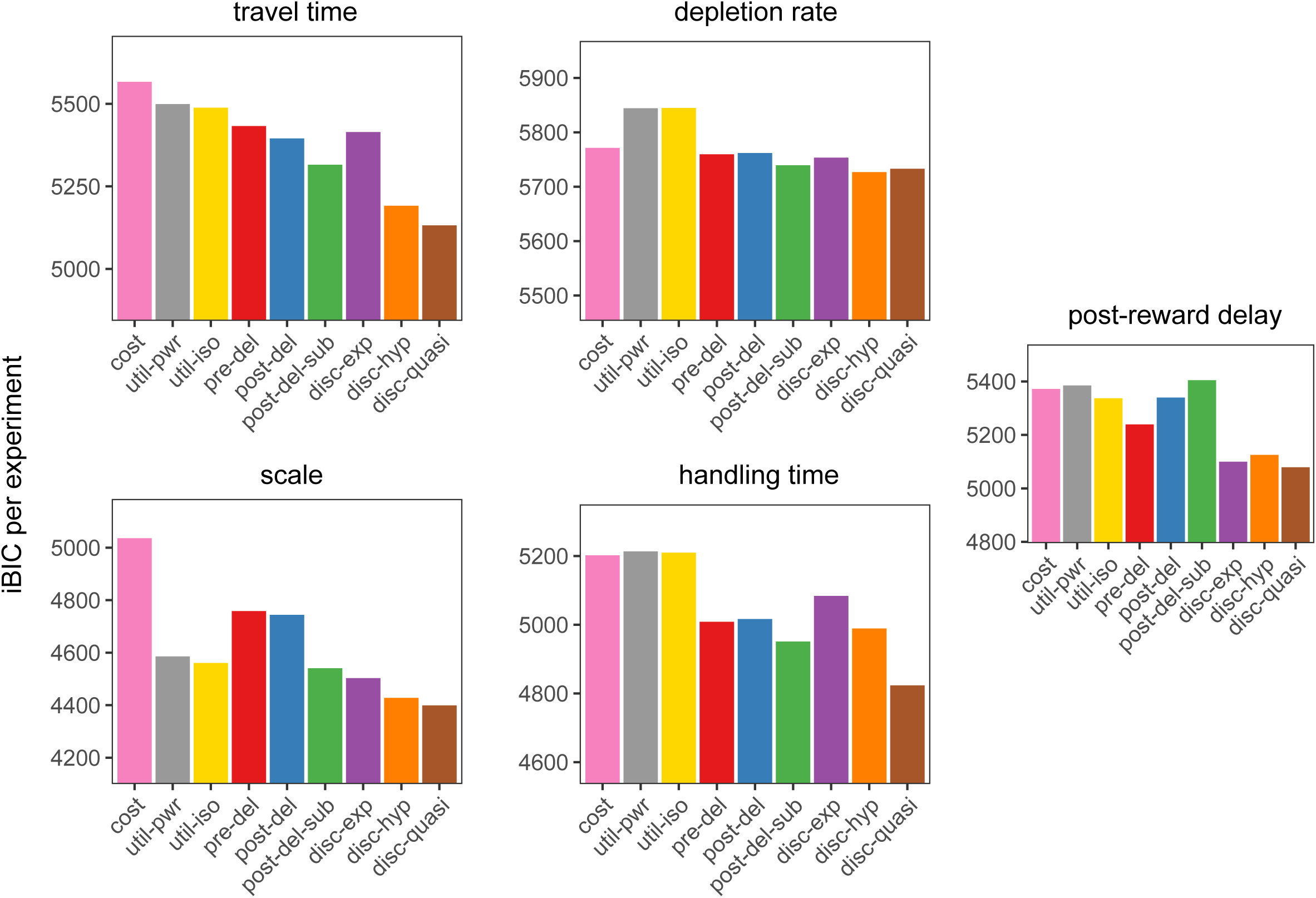
iBIC for each model for each foraging experiment.

**Figure S7.**
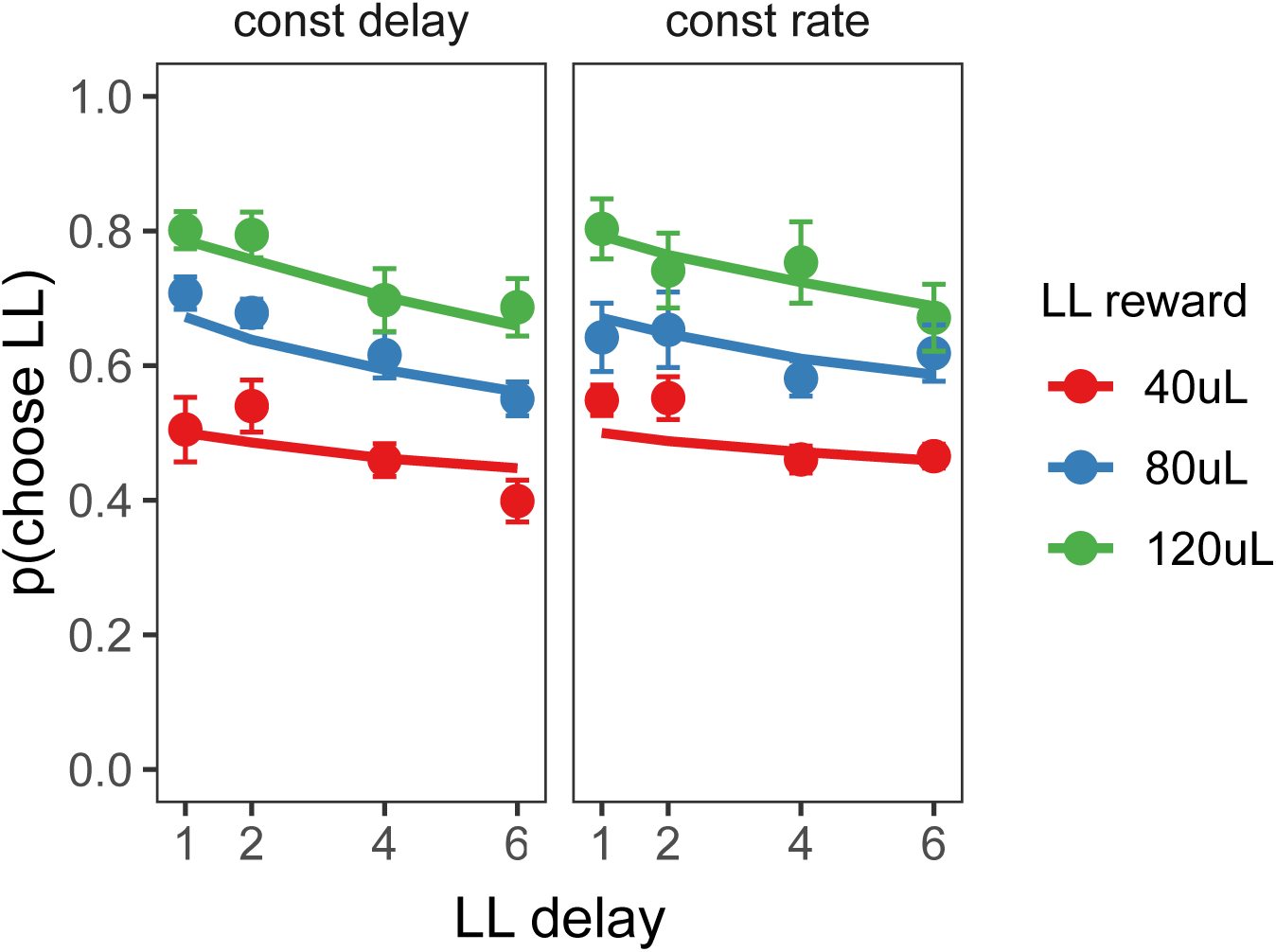
Predictions of the quasi-hyperbolic discounting model. Points and error bars represent mean ± standard error of rat behavior, lines represent mean quasi-hyperbolic model prediction.

**Figure S8.**
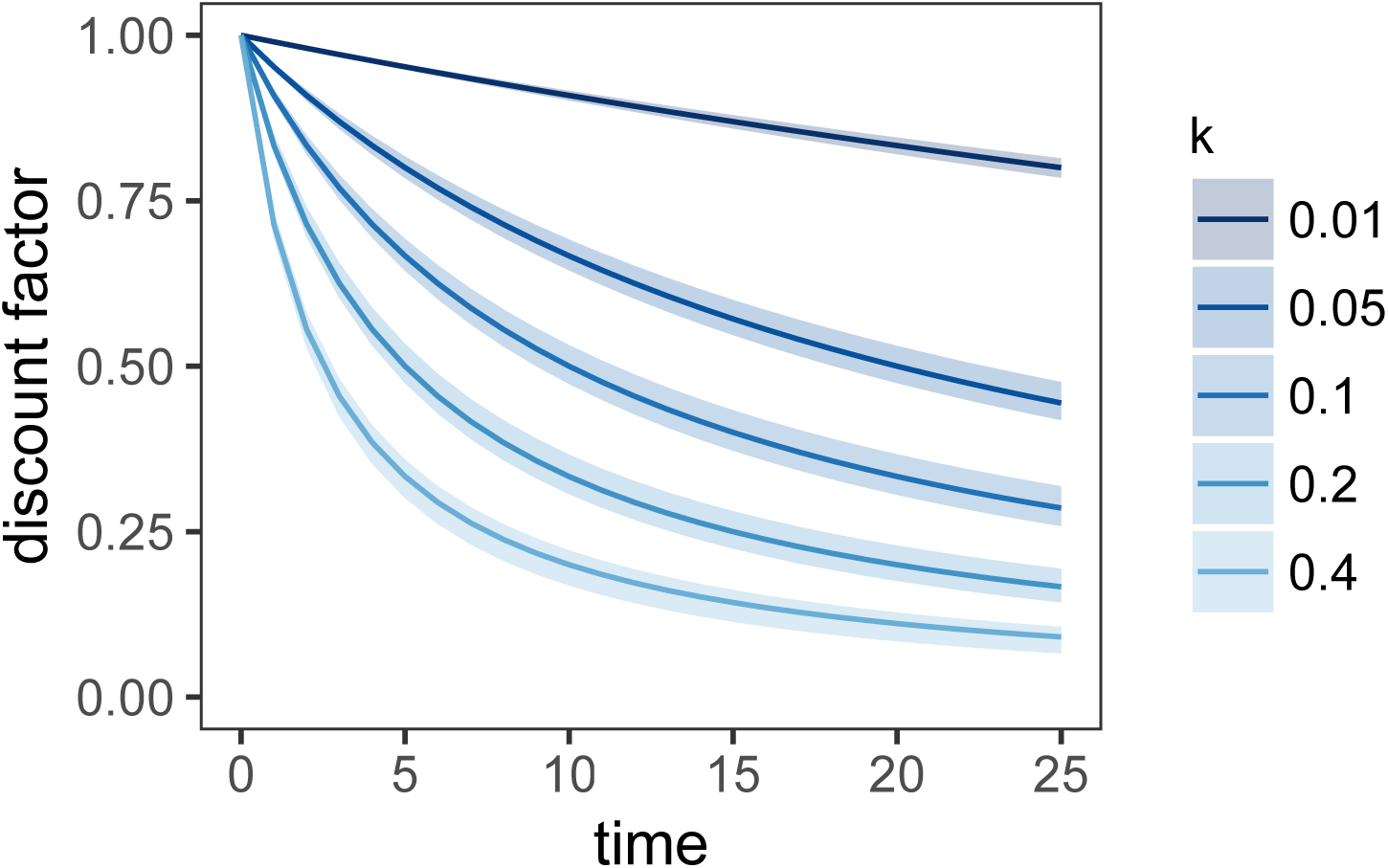
Discount function of the µAgent hyperbolic discounting model vs. standard hyperbolic discounting. Lines represent standard hyperbolic discounting function, 1/(1 + *k* * *time*). Ribbon represents the mean ± standard deviation of 100 simulations of the µAgent model in which the discount factor for each of the 100 µAgents was sampled from an exponential distribution with rate parameter *λ* = 1/*k*.

